# Calibration of a Closed-loop Model of Porcine Aortic Hemodynamics during Hemorrhage

**DOI:** 10.64898/2026.01.30.702699

**Authors:** Sadman Sadid, Matthew J. Eden, Fahim Usshihab Mobin, Micaela K. Gomez, Sandra Januszko, Heather Burkart, Lucas P. Neff, Timothy K. Williams, James E. Jordan, Elaheh Rahbar, C. Alberto Figueroa

## Abstract

Uncontrolled hemorrhage remains a leading cause of traumatic death, driven by rapid physiological deterioration that is often difficult to detect during the compensated phase. While large-animal models provide critical insights into these dynamics, they are resource-intensive, motivating the need for efficient computational frameworks that can mechanistically interpret cardiovascular responses. We developed and calibrated a closed-loop zero-dimensional (0D) lumped-parameter model (LPM) using hemodynamic data from 43 anesthetized swine subjected to controlled hemorrhage (10%, 20%, or 30% of total blood volume). The computational framework, incorporates a dynamic heart model with a custom time-varying elastance function, a multi-compartment aorta, and distal Windkessel models representing vascular beds. The model was calibrated at discrete time ‘snapshots’ throughout the 30-minute hemorrhage protocol to reproduce group-averaged experimental waveforms for aortic flow, regional organ flows, and systemic pressures. The calibrated model successfully reproduced experimental hemodynamic targets and waveform morphology across all hemorrhage severities. Analysis of the calibrated parameters revealed distinct physiological mechanisms driving hemodynamic adaptation during hemorrhage: a preferential increase in renal resistance compared to carotid resistance, indicating flow redistribution to vital organs, and a progressive mobilization of venous unstressed volume to sustain cardiac filling. Furthermore, the model captured the distinct shift toward preload limitation state for 30% hemorrhage group. This study establishes a physiologically interpretable in-silico framework capable of predicting both global and regional hemodynamic responses to acute blood loss, providing a validated foundation for future applications in trauma care and resuscitation modeling.

## 1 Introduction

Hemorrhagic shock is responsible for an estimated 1.5 million annual deaths worldwide and is the leading cause of preventable deaths under the age of 44 years [1]. In both civilian and military trauma, uncontrolled hemorrhage is a primary contributor to survivable fatalities, prompting the U.S. Department of Defense and Combat Casualty Care programs to identify hemorrhage control as a critical unmet need [2], [3], [4].

Hemorrhage is a rapidly evolving and time-critical physiological insult, with patient outcomes strongly determined by events occurring within the first hour following injury [1], [5]. Current trauma care prioritizes early recognition and prompt intervention during this narrow therapeutic window; however, acute blood loss triggers compensatory cardiovascular responses that can temporarily preserve arterial pressure and other vital signs despite substantial reductions in circulating volume, venous return, and cardiac output. As a result, physiological deterioration may remain clinically occult during the period when intervention is most effective. Once compensatory mechanisms fail, cardiovascular collapse can occur abruptly and may be difficult to reverse. A deeper understanding of the dynamic physiological response to hemorrhage is therefore essential for improving early assessment, interpreting hemodynamic signals, and guiding timely treatment during acute blood loss.

Direct investigation of hemorrhagic shock in humans is ethically constrained, often necessitating the use of analogs such as lower-body negative pressure [6], [7]. While these models replicate central hypovolemia, they do not fully capture the physiology of active blood loss [7]. Consequently, insight into the physiological response to hemorrhage has largely relied on large-animal models, with swine studies playing a central role due to close anatomical and cardiovascular similarity to humans [8], [9], [10], [11]. Controlled and uncontrolled hemorrhage experiments in swine facilitate careful monitoring of both the hemodynamic and physiological responses through invasive measurements and blood sampling. However, such studies are resource-intensive, requiring specialized surgical facilities, veterinary expertise, and significant financial investment and difficult to scale, motivating complementary approaches that can integrate and extend experimental observations.

Physiologically-based, experimentally-validated computational models provide a complementary framework to animal models for mechanistically interpreting cardiovascular responses to blood loss and for integrating sparse measurements across the circulation. Zero-dimensional (0D) lumped-parameter models (LPM) represent the circulation as interconnected, non-spatial compartments, analogous to an electrical circuit with elements such as resistors, capacitors, and inductors. Lumped-parameter models are well-suited for studying hemodynamic regulation during hemorrhage because of their computational efficiency, enabling them to simulate clinically relevant time scales (minutes to hours) in real time. Summers et al. validated an open-loop LPM computational platform against an human lower-body negative pressure data, demonstrating its ability to reproduce observed hemodynamic responses during simulated blood loss [12]. Curcio et al. analyzed and categorized multiple mathematical models of hemorrhagic shock, explicitly comparing their underlying physiological assumptions, and state variables used to represent hemorrhage-induced cardiovascular adaptation [13]. More detailed 0D formulations by Jin et al [14] and Beard et al [15] have incorporated mechanisms such as sympathetic stimulation and renin-angiotensin regulation to simulate blood pressure and heart rate responses during hemorrhage. Three-dimensional (3D) models enable high-fidelity representation of hemodynamics in image-based anatomical models. Renaldo et al. used 3D computational fluid dynamic techniques to study localized vascular responses under hemorrhage scenarios [16].

Despite advances in physiologically-based computational modeling of hemorrhage, several important limitations remain. Most 0D formulations have considered open-loop circuits and therefore have not provided a means of keeping track of blood volume, a key parameter during hemorrhage. Additionally, most 0D formulations have focused on reproducing global hemodynamic trends, such as mean arterial pressure (MAP) and cardiac output (CO), with limited attention to regional flow redistribution. Validating regional flow is critical because global metrics often remain stable during the compensated phase of shock, while significant differences in perfusion patterns between different vascular territories may exist. 3D models can provide valuable local details but are computationally infeasible for whole-body or long-duration hemorrhage simulations. Together, these limitations highlight the lack of models that can simultaneously remain computationally efficient, physiologically interpretable, and validated against detailed experimental hemorrhage data encompassing both global and regional hemodynamics.

To address this need, we developed and calibrated a closed-loop LPM using experimental data from 43 porcine subjects undergoing controlled hemorrhage (10%, 20%, and 30% total blood volume loss). The model reproduces left-ventricular pressure-volume loop dynamics, systemic pressure control, and branch-specific aortic flow distributions, at discrete analysis times (‘snapshots’) throughout hemorrhage. This model provides a first step towards the ultimate goal of developing a mechanistically interpretable in-silico framework to predict global and regional hemodynamic responses relevant to both civilian and military trauma care.

## 2. Methods

The methodological framework of this paper (**Figure 1**) is organized into three components: (i) description of the large animal hemorrhage model and experimental data acquisition; (ii) pre-processing of experimental data and generation of representative hemodynamic targets, and (iii) calibration of a reduced order (i.e., 0D) closed-loop computational model at discrete snapshots during hemorrhage.

**Figure 1:**
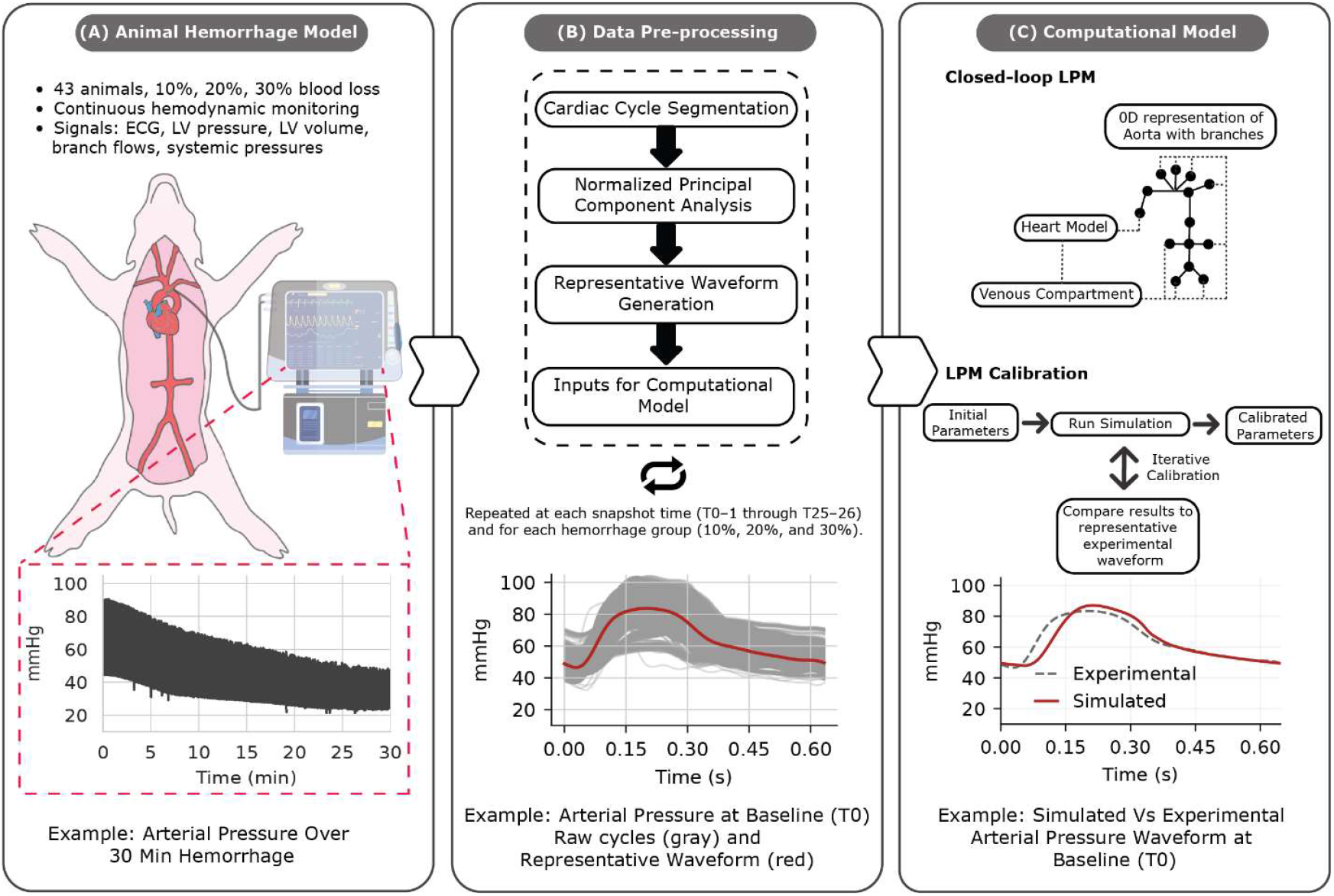
Overview of methods. (A) Controlled hemorrhage experiments in pigs with continuous hemodynamic monitoring. (B) Data preprocessing pipeline for representative waveform generation. (C) Closed-loop LPM and calibration process.

### 2.1 Animal Hemorrhage Model

All animal experiments were conducted under an approved protocol reviewed by the Wake Forest University Institutional Animal Care and Use Committee (IACUC). Adult Yorkshire swine weighing 55 to 80 kg (average 64 Kg) were split into three groups with an equal number of male and female subjects. Animals were anesthetized and instrumented with flow and pressure probes (Transonic Corporation, Ithaca, NY) at various locations in the aorta, a pressure-volume catheter in the left ventricle (Transonic Systems Inc., Ithaca, NY, USA), and a central venous pressure catheter (**Figure 2**). Either 10% (N =15), 20% (N = 15), or 30% (N = 13) of the blood volume was removed at a constant rate through the left femoral vein over a 30-minute period, assuming an initial blood volume of 60 ml/kg. For the present analysis, only the first 25 minutes of the hemorrhage phase were used, as data beyond that point corresponded to a subsequent resuscitation intervention during which a REBOA catheter was introduced.

**Figure 2:**
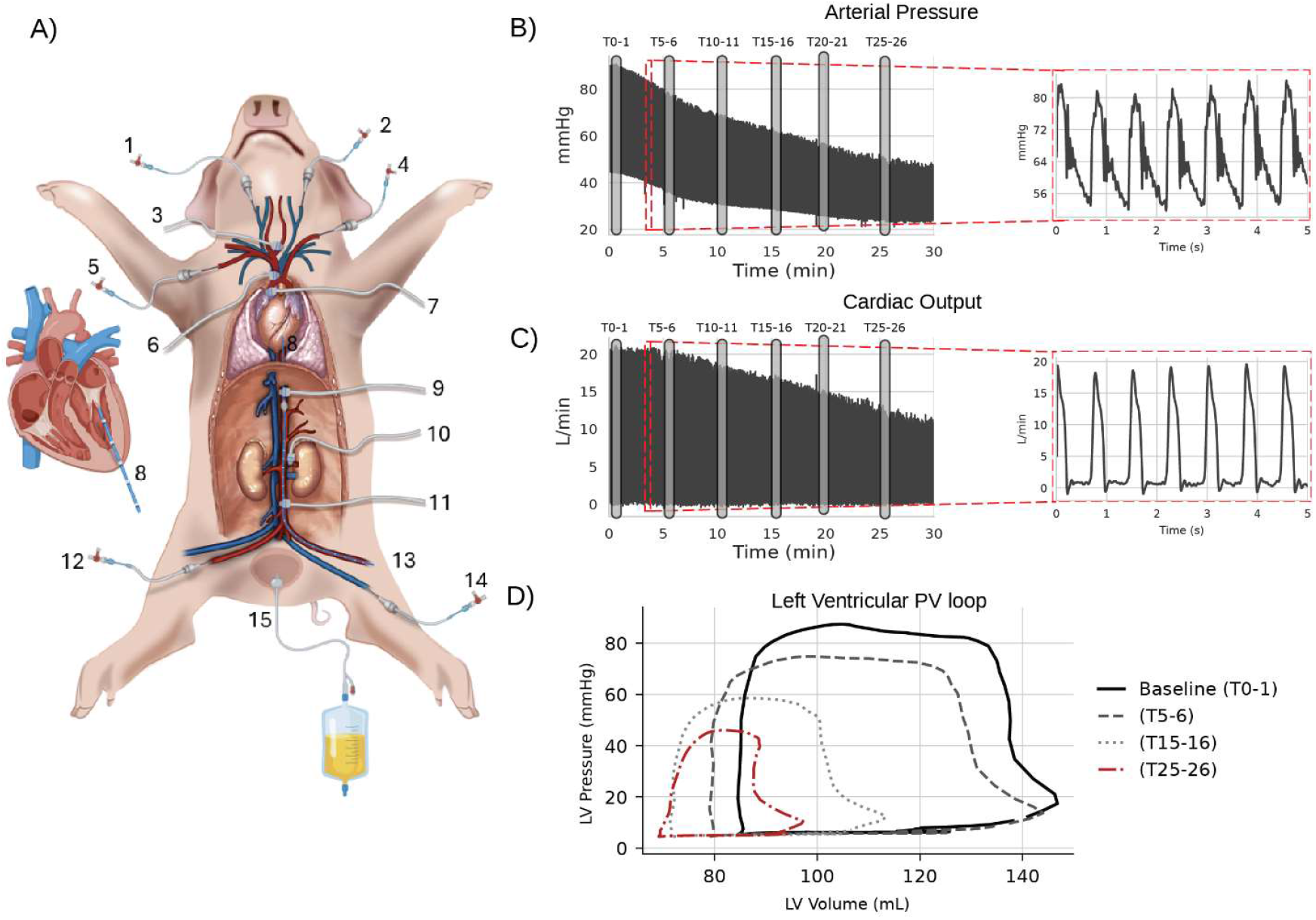
(A) Diagram of the location of the various flow and pressure probes. **Flow Probes**: (3) Left Carotid Artery; (6) Brachiocephalic arterial trunk; (7) Ascending Aorta; (9) Descending Aorta; (10) Left Renal Artery; (11) Infra-renal Aorta. **Pressure Probes**:(4) Left Subclavian Artery; (12) Right Femoral Artery, measuring pressure at the infra-renal aorta. **Left Ventricular PV catheter**: (8). **Hemorrhage Location**: (14) Left Femoral Vein. **Additional Probes:** (1) Right External Jugular Vein central venous pressure (CVP) monitoring; (2) Left External Jugular vein, fluid and drug administration catheter; (5) Right Brachial Artery, blood sampling catheter; (15) Urine collection catheter; (13) Left Femoral Artery, 9 Fr. cannula for REBOA balloon catheter (not used in this study). (B)Time history of Arterial Pressure - location (4), for a single individual. (C) Time history of Cardiac Output – location (7). (D) PV loop traces - location (8).

### 2.2 Data Pre-processing

Experimental hemorrhagic data including ECG, left ventricular pressure *LVP* and volume *LVV*, and multiple arterial pressure *P* and flows *Q* (**Figure 2**) were continuously collected over the course of 25 min. Since our goal is to calibrate a closed-loop 0D model of hemodynamics at discrete analysis times (‘snapshots’) throughout hemorrhage, we must first define representative hemodynamics at such snapshots describing all subjects included in each hemorrhage group. Towards that goal, six one-minute intervals were identified, separated by five minutes: T0-1, T5-6, T10-11, T15-16, T20-21 and T25-26. Each one-minute interval contains many cardiac cycles with some cycle-to-cycle variability for each subject. In this work, we assumed that the hemodynamic alterations due to hemorrhage are small within each one-minute interval, and that we can therefore define a single ‘snapshot’ (i.e., representative periodic waveform) for each interval:

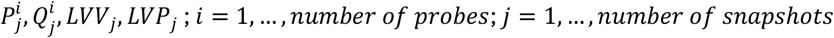

where *i* is an index describing each probe location, and *j* is the snapshot index. Ultimately, each snapshot describes the typical group level behavior observed during the corresponding time window, providing a time and population averaged characterization.

We implemented a two-step preprocessing workflow to construct these snapshot representative waveforms. First individual waveforms over the one-minute interval were isolated by defining the onset of systole for each cardiac cycle (Section 2.2.1). Second, representative waveforms were generated for each signal at each snapshot time for each hemorrhage group (Section 2.2.2) using principal component analysis PCA. Finally, to complete the data processing, left ventricular elastance functions were derived using the representative LV pressure and volume waveforms.

#### 2.2.1. Estimating the Onset of Systole from LV Pressure Using dLVP/dt

The ECG R-wave is commonly used to identify the start of systole. However, only 24 of 43 subjects had ECG recordings with reliable R-wave identification over the full hemorrhage protocol. In contrast, high quality LVP signals were available for all animals. Therefore, we used a the subjects with both reliable R-wave ECGs and LVP recordings to develop an algorithm for estimating systolic onset directly from LVP to define the heart rate. In this approach, systolic timing was defined using the maximum of the first derivative of LVP dLVP/dt. Although the timing of maximum *dLVP*/dt does not coincide exactly with the ECG R-wave peaks [17], [18], it provides a reliable temporal landmark across animals, enabling robust temporal alignment and extension of this approach to subjects with unreliable ECG waveforms.

This algorithm was defined as follows. First, for subjects with reliable ECG signals and clearly detectable R-waves (**Figure 3A**), R-peak locations were identified. We then used the corresponding LVP waveform to compute dLVP/dt and detect successive maxima(**Figure 3B**). For each beat, the interval between two consecutive dLVP/dt maxima was defined as T_LVP_, while the interval from a dLVP/dt maximum to following ECG R-peak was defined as T_R-wave_ (**Figure 3B**). The ratio of these two intervals, defined as R-peak fraction = T_R-wave_/T_LVP_, was then used to define systolic onset from LVP data for subjects with usable ECG waveforms.

**Figure 3:**
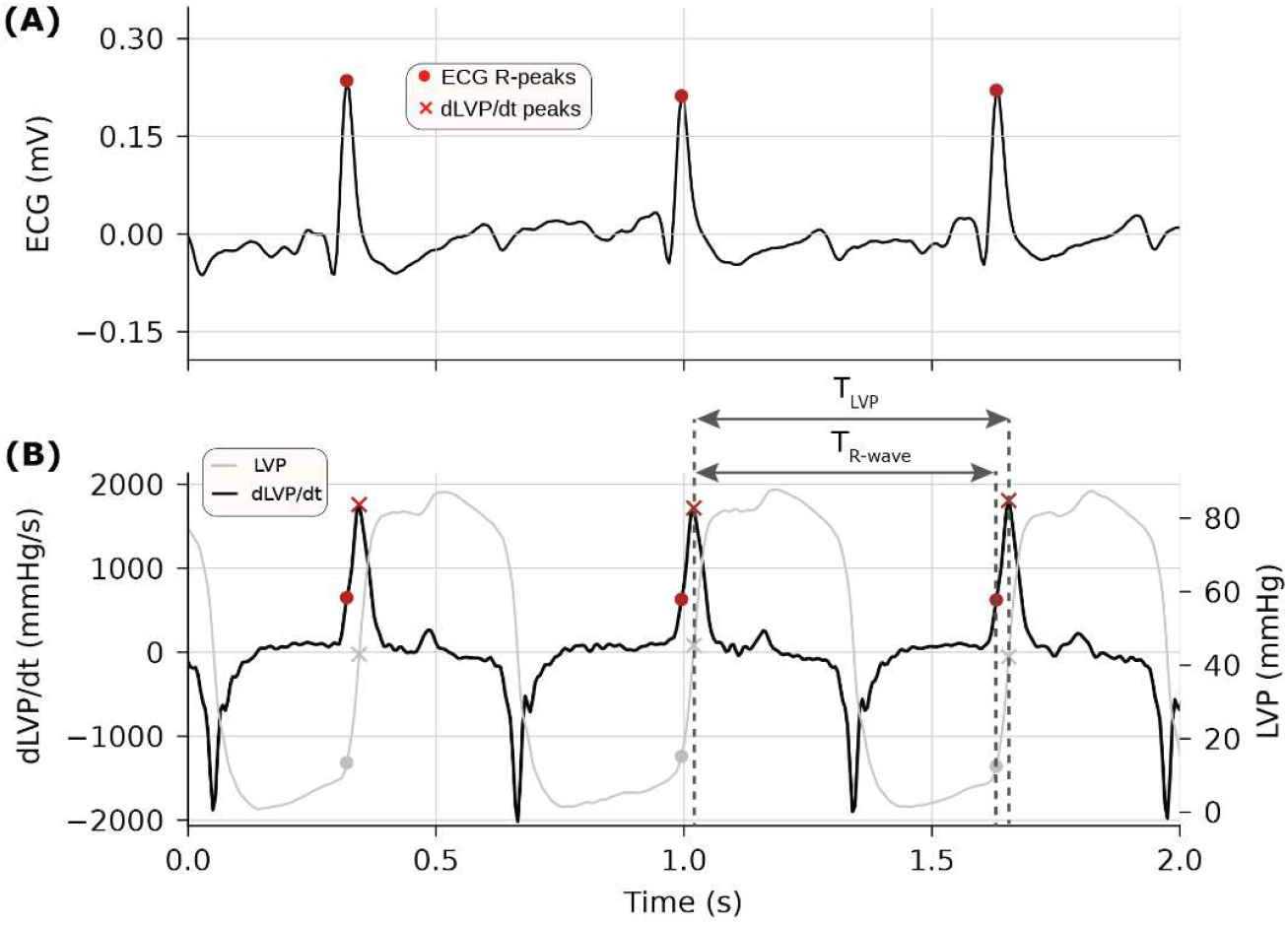
Estimating systolic onset from LV pressure using dP/dt. Single-subject example at T0-1. **(A)** ECG waveform with detected R-peaks (red ●). **(B)** Left-ventricular pressure (LVP, gray) and its time derivative dLVP/dt (black), with dLVP/dt maxima (red ×). The interval **a** denotes the duration between two consecutive dLVP/dt maxima, while **b** denotes the relative position of the ECG R-peak within this interval. This fractional timing is used to estimate systolic onset from LV pressure alone.

R-peak fraction variability with hemorrhage was quantified for each group. R-peak fractions were computed for each cardiac cycle within each one-minute interval, then pooled across subjects with reliable ECG within each hemorrhage group and interval. Median and interquartile range (IQR) for R-peak fractions was quantified (**Figure 4A**). Interestingly, the R-peak fraction decreased over time with hemorrhage and was smallest for the 30% hemorrhage group, indicating a separation between R-waves and maximum dLVP/dt with increasing levels of blood loss. The calculated R-peak fractions were then used to estimate systolic onset directly from LVP for subjects lacking reliable ECG signals within a group. Heart rates could then be computed and summarized for all subjects within each hemorrhage group over the one-minute intervals (median and IQR; **Figure 4B**).

**Figure 4:**
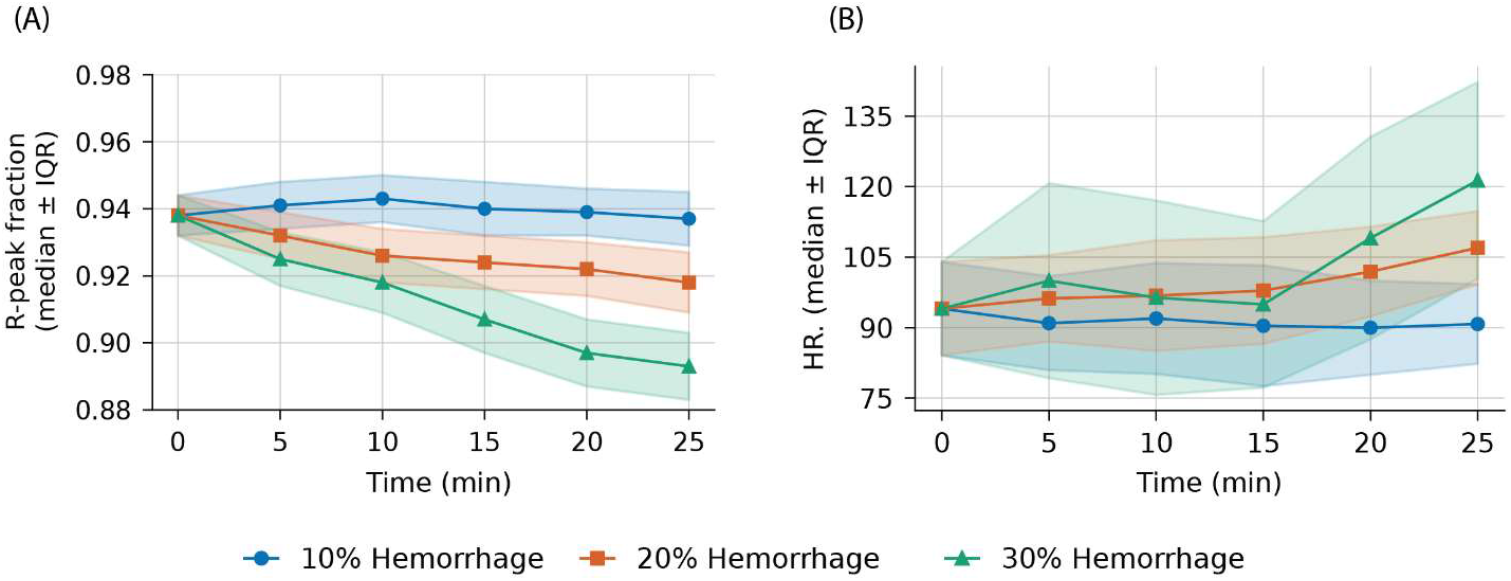
(A) Time course of the median R-peak fraction within the dLVP/dt-defined cardiac cycle for the three hemorrhage groups, computed over successive one-minute intervals from baseline through 25 min. (B) Corresponding heart-rate trajectories over the same intervals. Points indicate group medians; shaded regions denote interquartile ranges (IQR).

#### 2.2.2 Constructing Representative Hemodynamic Waveforms Using PCA

Using the above method, we defined cardiac cycle lengths for each subject in each hemorrhage group and for each one-minute interval. All waveforms within each one-minute interval were then condensed into a group-specific representative waveform (snapshot) that captures the typical cardiac-cycle pattern, including both timing and amplitude, across subjects despite substantial intra- and inter-animal variability. To achieve this, a two-stage approach was employed. First, principal component analysis PCA was used to extract a representative normalized (time and amplitude) waveform capturing the dominant shape features. Second, the normalized waveform was rescaled back to physical units based on the group level signal amplitude.

##### Stage 1: Normalization and PCA for shape extraction

For a fixed hemorrhage group *G* ∈ [10%, 20%, 30%] and a given one-minute interval *j*, let *x*_*n*_ (*t*) be the *n*^th^ waveform for a measured signal for a given animal within the group *G*. This waveform is segmented and resampled to a uniform grid of *M* phase points by cubic interpolation. To isolate shape variability, min-max amplitude normalization was performed, producing a normalized vector for each waveform *n*^th^ 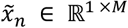. All normalized waveforms were then concatenated into a data matrix *X* ∈ ℝ^*N* ×*M*^, where *N* is the total number of waveforms within the one-minute for all subjects in the hemorrhage group *G*.

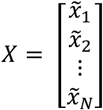

Principal component analysis (PCA) was applied to the matrix *X* to reduce dimensionality and to extract dominant modes of shape variation [19]. For each signal (*P*^*i*^, *Q*^*i*^, *LVV, LVP*), a representative normalized waveform *y*_*G,j*_ defining the snapshot *j* was calculated as:

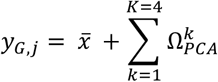

Where 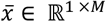 is the mean waveform of *X*, and 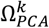 represent each of the first four PCA modes, which together explained most of the waveforms variance. This reconstruction captures the dominant waveform morphology for the entire group while reducing sensitivity to outlier beats. The resulting waveform provides a low-dimensional description of shape; independent of amplitude (**Figure 5A, 5C**).

**Figure 5:**
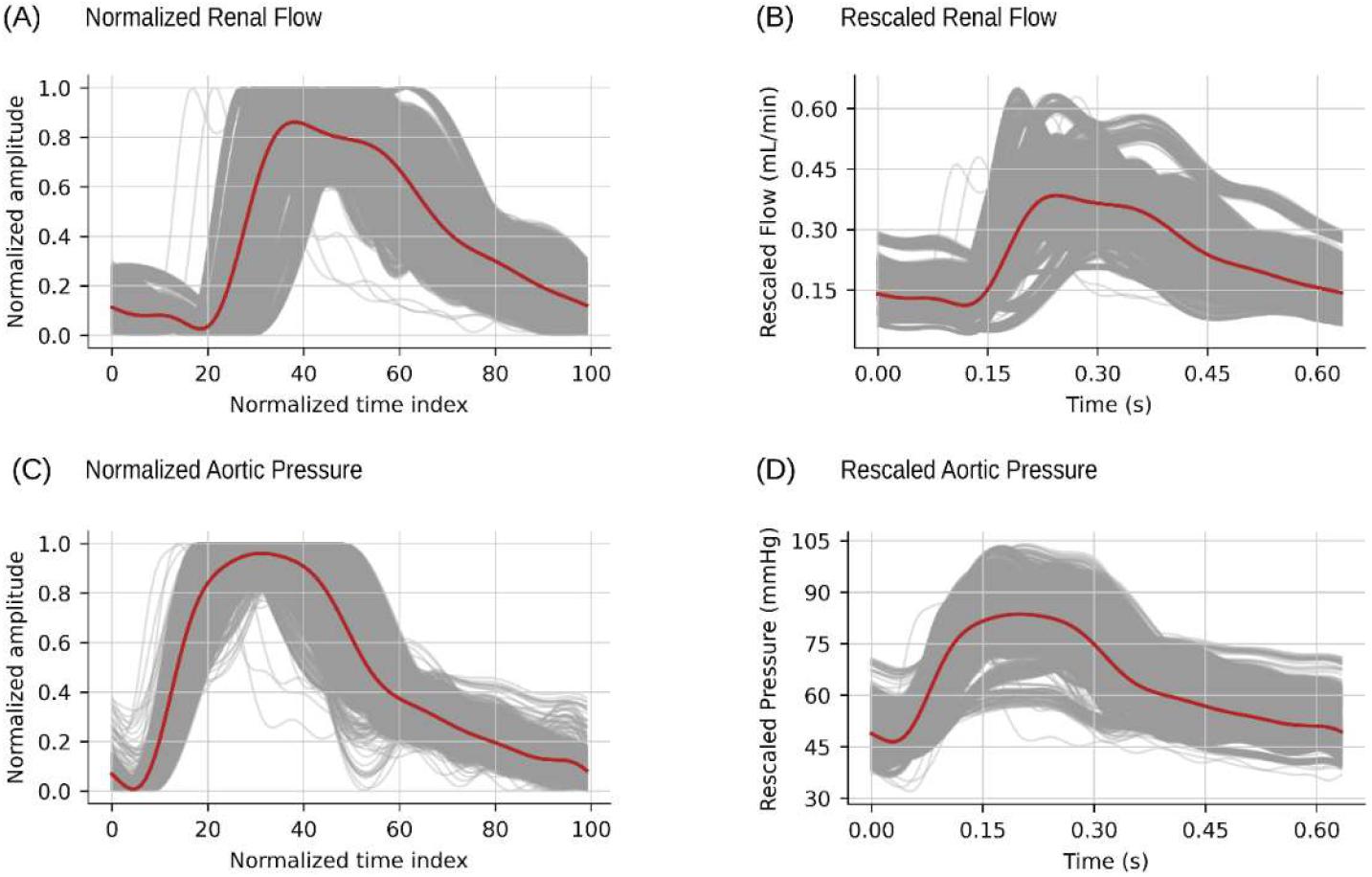
(A), (C) Normalized renal flow and aortic pressure waveforms at the baseline T0-1 interval (gray) with the representative PCA reconstruction (red). Rescaled waveforms defining the representative snapshot hemodynamics are shown in panels (B)(D).

##### Stage 2: Rescaling back to physiological units

The normalized group representative waveforms obtained in stage 1 capture the overall morphology of each signal (*P*^*i*^, *Q*^*i*^, *LVV, LVP*) but are normalized both in time and amplitude. To make the waveforms physiologically interpretable, they must be rescaled back into physiological units. Towards that goal, we defined a common baseline reference (T0-1) across all subjects (N=43) regardless of the hemorrhage group *G*.

For each subject *s*, the mean *μ* and standard deviation *σ* for each recorded signal at baseline and each subsequent one-minute interval *j* were computed as, (*μ*_*s*,o_, *σ*_*s*,O_) and (*μ*_*s,j*_, *σ*_*s,j*_) respectively.

Subject-level ratios at interval *j* for both the mean and the standard deviation describing changes relative to baseline were defined as,

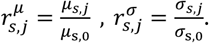

Group-level (*G*) scaling factors were then obtained as the median across subject level ratios for all subjects in a particular group (*s* ∈ *G*):

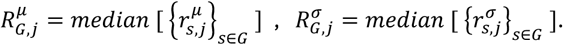

For group *G* at interval *j*, the mean and standard deviation for each of the recorded signals were then computed as:

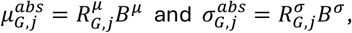

where, *B*^*μ*^ and *B*^*σ*^ denote the median mean and median standard deviation at baseline calculated across all subjects:

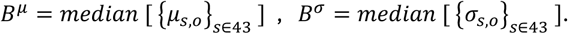

Each normalized PCA representative waveform *y*_*G,j*_ (with, mean 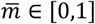 and standard deviation *std* ∈ [0,1]) was then rescaled back to a physiological amplitude waveform 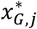 as,

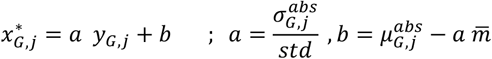

Lastly, each normalized phase point *M* was mapped back to physical time *t* using the median cardiac cycle duration across the hemorrhage group *G* for all intervals *j*, yielding a consistent temporal reference for comparison across groups. The resulting waveform, 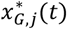 restores both the time and physiological amplitude for the whole group *G* and any interval *j* for all the recorded signals (*P*^*i*^, *Q*^*i*^, *LVV, LVP*). Examples of the rescaled waveforms are shown in **Figure 5B** and **5D**.

#### 2.2.3 Deriving Representative Left Ventricular Elastance Waveform

An instantaneous left ventricular elastance 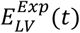 for each hemorrhage group *G* and interval *j* was computed from the snapshot *LVV, LVP* data as;

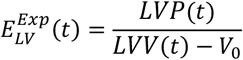

where *V*_O_ represents the left ventricular unstressed volume. A constant value of *V*_O_ = 17.8 *mL* was used, consistent with previously reported porcine measurements [20]. This left ventricular elastance 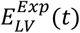 will be used to inform an analytical elastance function *E*_*LV*_(*t*)which drives a heart model in the closed-loop LPM.

### 2.3 Computational Model

A closed-loop LPM was defined using the open-source cardiovascular simulation software CRIMSON [21]. A key objective of this work was to develop and calibrate a computational model that reproduces experimental hemodynamics under progressive hemorrhage. This closed-loop model includes different LPMs such as a heart model with time-varying elastance function, an aortic model, a series of 3-element Windkessel models representing distal vascular beds in the arterial circulation, and a venous compartment model (see **Figure 6**). The following section describes in detail the different LPMs, as well as the calibration process of all model parameters in detail.

**Figure 6:**
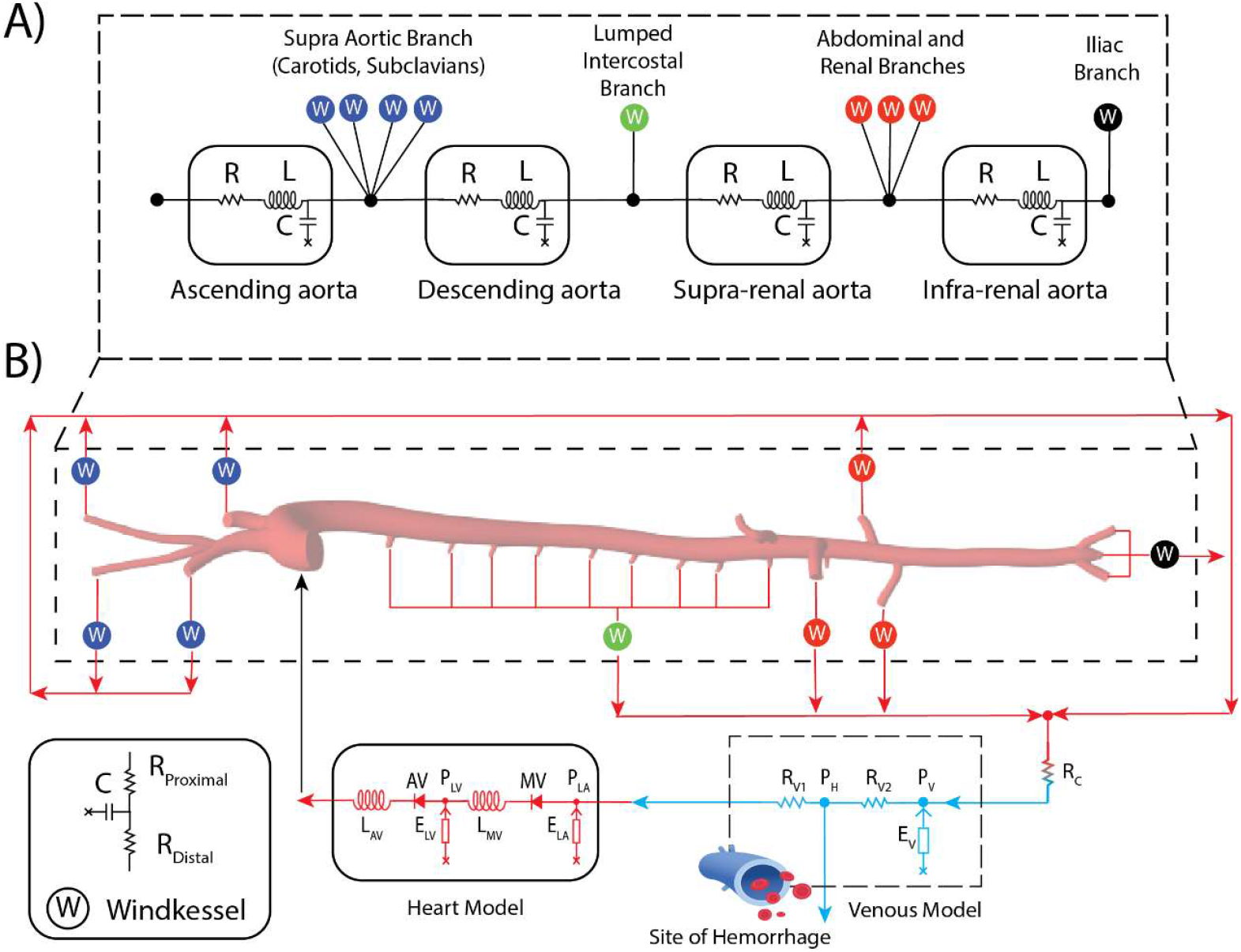
A) Lumped parameter representation of the Aorta B) in-silico closed-loop lumped parameter network.

#### 2.3.1 Closed-loop LPM

##### (1) Lumped parameter Heart model

Rather than prescribing inlet pressure or flow waveforms directly, we employed a dynamic lumped-parameter heart model to generate physiologically consistent pulsatile flow waveforms in response to time-varying loading conditions. The lumped-parameter heart model consists of a left atrium, mitral valve, left ventricle, and aortic valve (**Figure 6B**). The pumping action of the left ventricle was modeled with a pressure-generating chamber driven by a time-varying elastance function *E*_*LV*_ (*t*), while the left atrium was represented as a fixed-compliance chamber *E*_*LA*_ that passively fills and empties in response to transvalvular pressure gradients. The mitral and aortic valves were modeled as perfect (e.g., non-regurgitant) diodes with small inertances in series, ensuring unidirectional forward flow during valve.

##### Analytical Elastance Function

Previous studies [22], [23] have reported that porcine left ventricular elastance waveforms exhibit four distinct characteristic segments (see **Figure 7A**) representing a rapid systolic upstroke (segment I), a slower systolic upstroke (segment II), followed by a rapid diastolic decay (segment III) and a constant diastole phase (segment IV). These segments are also evident in our experimental elastance waveforms 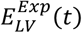. These segments are separated by markers t_mid_, t_max_, and t_min_.

**Figure 7:**
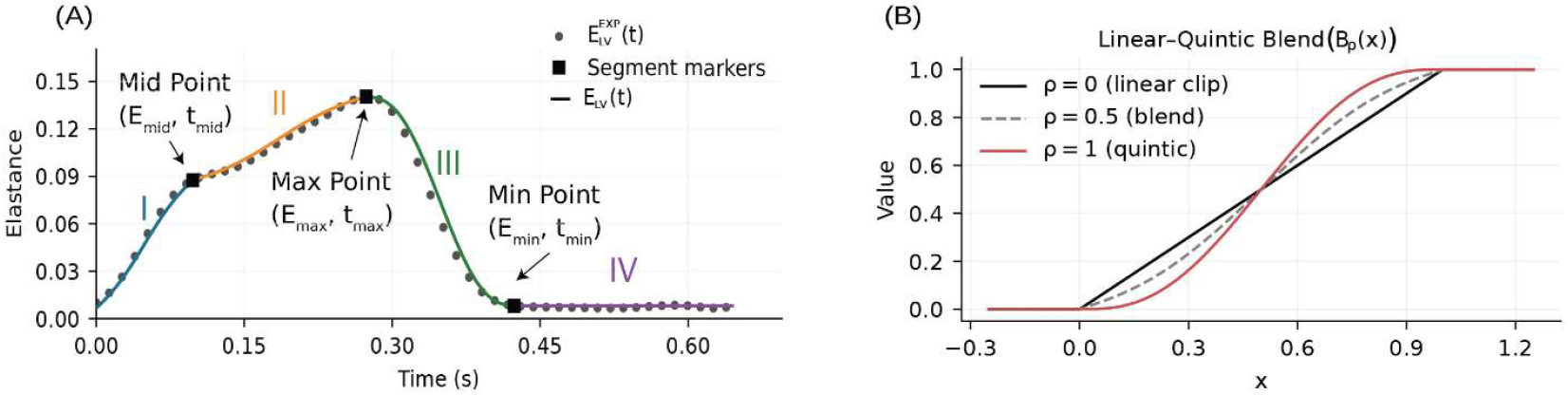
Analytical left ventricular elastance *E*_*LV*_(*t*) (solid line) at the baseline (T0−1, N=43). (A) Experimental elastance waveform 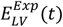 (black dots) and fitted analytical elastance function *E*_*LV*_(*t*) (solid line). Segment markers separate four distinct segments: rapid systolic upstroke (I), slower systolic upstroke (II), rapid diastolic decay (III), and near constant diastolic phase (IV). (B) Linear−quintic blending function *B*_*ρ*_(*x*) illustrating the effect of the curvature parameter *ρ*, which controls the smoothness of transitions between elastance segments, ranging from a purely linear clip (*ρ* = 0) to a fully quintic smoothstep (*ρ* = 1).

Previous broadly used left ventricular elastance analytical functions included a sinusoidal model by by Pope [24], and a Double-Hill model by Mynard [25]. However, these models fail to capture the four distinct segments experimentally observed. Towards that goal, in this paper we developed a custom analytical elastance function. This custom function combines two main components, a quintic *smoothstep f*unction [*S*(*x*) = 6*x*^5^ − 15*x*^4^ + 10*x*^3^] [26] as well as a *linear clip* function [*clip*(*x*, 0,1)]. These two components are blended into a transition operator (*B*_*ρ*_(*x*)):

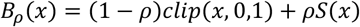

*ρ* ∈ [0,1] is a parameter controlling the degree of slope and curvature: *ρ* = 0 yields a purely linear ramp and *ρ* = 1 yields a quintic-shaped sigmoidal transition (**Figure 7B**). Using this transition operator, the analytical time varying elastance *E*_LV_ (*t*) was then defined piecewise over the four characteristic segments as:

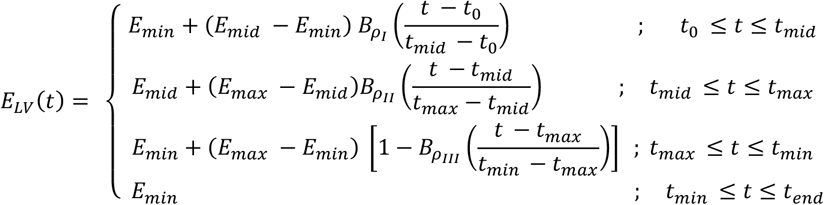

Here, *E*_*min*_, *E*_*mid*_, *E*_*max*_ represent the experimental elastance values at diastole, systolic shoulder inflection point, and peak systole, respectively. *E*_*min*_ and *E*_*max*_ were directly obtained from the minimum and maximum of the experimental elastance waveform, while *E*_*mid*_ was identified as the inflection point where the rapid systolic upstroke transitions into the slower systolic upstroke stage. *t*_O_, *t*_*end*_ represent the start and end of the cardiac cycle. The curvature parameters (*ρ* _*I*_, *ρ* _*II*_, *ρ* _*III*_) control the shape of each segment and define the corresponding transition operators *Bρ* _*I*_,*Bρ* _*II*_, *Bρ* _*III*_. The curvature parameters were estimated using the procedure described in the Elastance Model Parameter Calibration section below.

##### (2) Aortic model

The porcine aorta was represented as LPM composed of four sequential compartments: ascending aorta, descending aorta, supra-renal aorta, and infra-renal aorta (**Figure 6A**). Each compartment was modeled as a three-element RLC unit composed of resistance R, inertance L, and compliance C to capture viscous loss, blood inertia, and elastic storage, respectively [27], [28].

##### (3) Distal Arterial Windkessel Models

To characterize the systemic circulation downstream of the aortic compartments, distal arterial branches were represented as three-element Windkessel (RCR) models at their corresponding aortic outlets (**Figure 6B**). Each Windkessel model consists of a proximal resistance *R*_*p*_, compliance *C* and distal resistance *R*_*d*_, capturing the impedance of downstream vascular beds. To reduce the number of model parameters, certain branches were aggregated: the intercostals into a single lumped intercostals outlet, the celiac and superior mesenteric arteries were into a single abdominal outlet and the iliac arteries into a single iliac outlet.

##### (4) Venous Compartment

Distal resistance *R*_*d*_ from all the arterial Windkessel model outlets drained into a single venous compartment through a terminal resistance *R*_*c*_ representing the capillary beds. The venous compartment consists of two resistances *R*_*v*1_, *R*_*v*2_ and an elastance *E*_*v*_, representing the blood reservoir behavior of the venous system (**Figure 6B**). The venous elastance relates pressure at node *P*_*v*_ with total circulating blood volume *V*_*v*_ and unstressed volume *V*_*u*_ as follows:

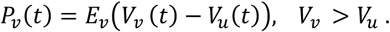

The venous unstressed volume *V*_*u*_ defines the portion of venous blood that does not contribute to active pressure generation [29]. Changes in *V*_*u*_ therefore alter circulating volume and impact venous return and cardiac filling.

##### Hemorrhage model

Hemorrhage was modeled by progressively reducing the total circulating blood volume *V*_*v*_ available within the closed-loop circuit at each hemodynamic snapshot *j*. The total pre hemorrhage (T0-1) circulating volume was estimated as *V*_*v* TO−1_ = 60 *mL*/*kg* [20].

Each hemorrhage group *G* was defined by a target hemorrhage fraction *H*_*G*_ ∈ [10%, 20%, 30%] of total blood loss over a 30-minute period. Therefore, for each hemodynamic snapshot *j* and for each group G, the circulating volume was computed as,

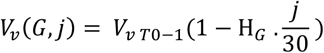

This volume was assigned to the venous compartment for all hemodynamic snapshots *j*. Prescribing the history of total circulating volume in this manner enables hemorrhage-induced reductions in venous return, preload, and systemic pressure to emerge naturally from the closed-loop pressure− volume relationships. The unstressed volume *V*_*u*_ will be calibrated for each snapshot according to the procedure described in the following section.

#### 2.3.2. LPM Calibration

LPM calibration was performed to reproduce experimental hemodynamics for each snapshot *j*. Left ventricular elastance and vascular model LPMs (i.e., aortic model, distal arterial Windkessel models, and venous compartment model) parameters were calibrated separately. The details of elastance and vascular LPM calibration at baseline and during hemorrhage are presented below.

### Elastance Model Parameter Calibration

The analytical elastance function *E*_*LV*_(*t*) contains nine parameters: three elastance level values, three corresponding timing parameters, and three curvature parameters. The elastances *E*_*min*_, *E*_*mid*_, *E*_*max*_ and corresponding timings *t*_*min*_, *t*_*mid*_, *t*_*max*_ were directly extracted from the experimental 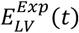for each hemodynamic snapshot as described earlier. The curvature parameters (*ρ*_*I*_, *ρ*_*II*_, *ρ*_*III*_) controlling the shape of the transition between segments in *E*_*LV*_ were calibrated using nonlinear least-squares minimization:

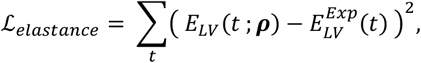

where ***ρ*** = (*ρ*_*I*_, *ρ* _*II*_, *ρ* _*III*_). The resulting calibrated analytical elastance functions were then prescribed as inputs to the lumped-parameter heart model for all snapshot simulations.

### Vascular Model Parameter Calibration

#### Overall strategy

The goal of the Vascular Model parameter calibration was to identify a set of model parameters (aortic model, distal arterial Windkessel models, and venous compartment model) that minimize discrepancies between close-loop model predictions and experimental pressure and flow data at each snapshot. For any given snapshot, initial guesses for LPM of vascular model were defined using literature data [30], [31], [32], and the following cost function was minimized using a local, derivative free optimizer (Powell method [33], [34]):

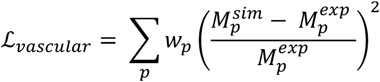

where *p* = 1, …, number of hemodynamic targets (described below) included in calibration. 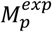 denotes *p*-th experimental target metric and 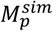 denotes the corresponding simulated metric, and *w*_*p*_ denote the weights for the different targets. Given the smaller number of pressure measurements, a larger weight (4.5x) was given to these compared to the flow targets.

The full set of LPM parameters of the vascular model was calibrated at T0-1. Baseline calibration variables included: Aortic RLC parameters, Windkessel parameters (*R*_*p*_, *R*_*d*_, *C*), capillary resistance *R*_*c*_, venous resistances *R*_*V*1_, *R*_*V*2_, venous compliance *E*_*V*_, and venous unstressed volume *V*_*u*_ as well as left atrial elastance *E*_*LA*_. For the subsequent hemorrhage snapshots, only a subset of vascular model parameters were further calibrated relative to their T0-1 value. These include Windkessel parameters and the venous unstressed volume *V*_*u*_, see Table 1. This assumes that the hemodynamic parameters of aorta, capillary beds, and venous compartment do not change during hemorrhage. All hemodynamic changes in the vascular model during hemorrhage are therefore capture by adjustments in the Windkessel parameters and *V*_*u*_. This approach has been previously adopted by others [31], [32], [35].

**Table 1:**
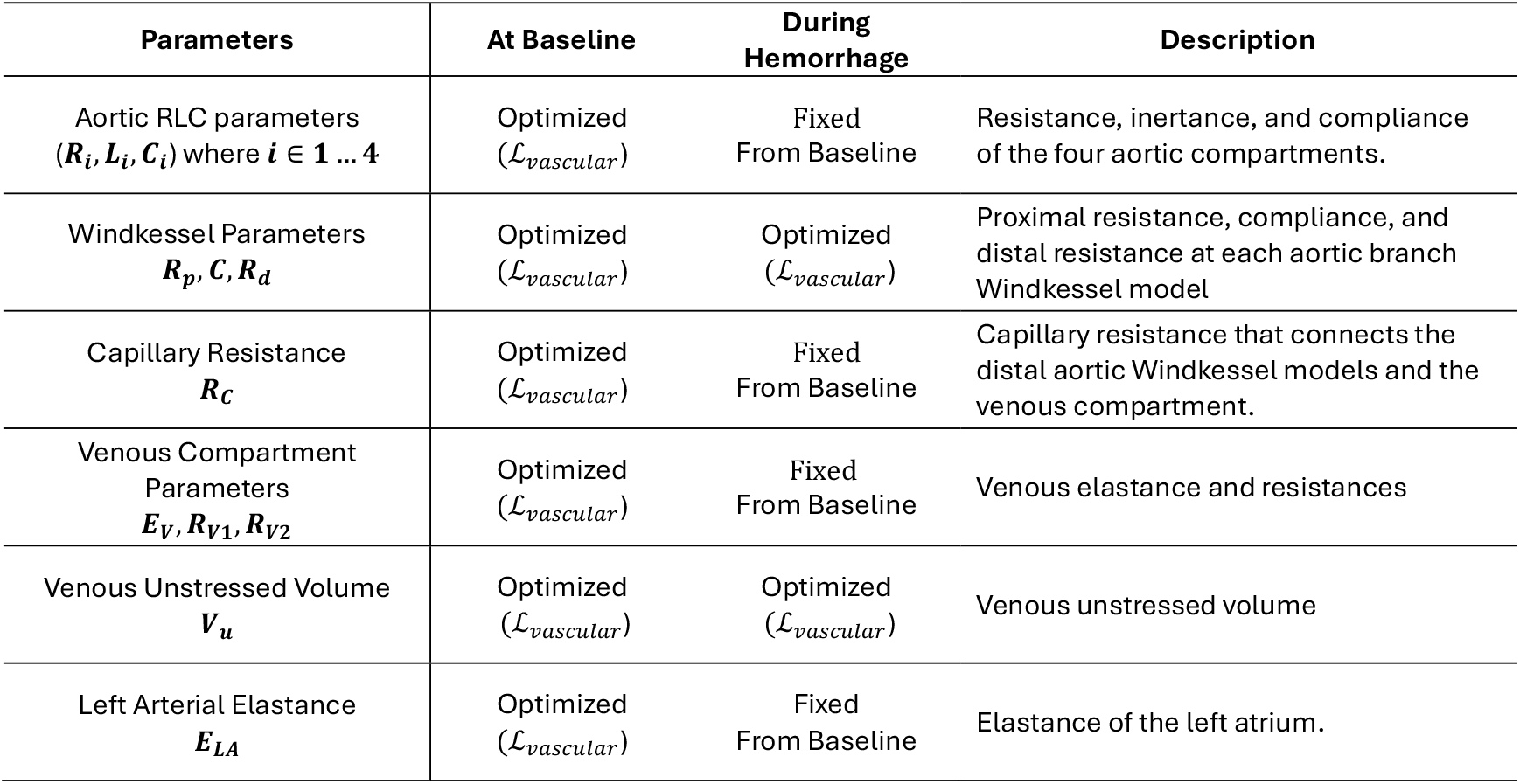
Summary of model parameters calibrated at baseline and during hemorrhage.

| Parameters | At Baseline | During Hemorrhage | Description |
| --- | --- | --- | --- |
| Aortic RLC parameters<br>($R_i, L_i, C_i$) where $i \in 1 \dots 4$ | Optimized<br>($\mathcal{L}_{vascular}$) | Fixed<br>From Baseline | Resistance, inertance, and compliance of the four aortic compartments. |
| Windkessel Parameters<br>$R_p, C, R_d$ | Optimized<br>($\mathcal{L}_{vascular}$) | Optimized<br>($\mathcal{L}_{vascular}$) | Proximal resistance, compliance, and distal resistance at each aortic branch Windkessel model |
| Capillary Resistance<br>$R_c$ | Optimized<br>($\mathcal{L}_{vascular}$) | Fixed<br>From Baseline | Capillary resistance that connects the distal aortic Windkessel models and the venous compartment. |
| Venous Compartment Parameters<br>$E_v, R_{v1}, R_{v2}$ | Optimized<br>($\mathcal{L}_{vascular}$) | Fixed<br>From Baseline | Venous elastance and resistances |
| Venous Unstressed Volume<br>$V_u$ | Optimized<br>($\mathcal{L}_{vascular}$) | Optimized<br>($\mathcal{L}_{vascular}$) | Venous unstressed volume |
| Left Arterial Elastance<br>$E_{LA}$ | Optimized<br>($\mathcal{L}_{vascular}$) | Fixed<br>From Baseline | Elastance of the left atrium. |

#### Flow and Pressure Targets

Flow measurements were available at (see **Figure 8**) Left Carotid Artery (LCA); Brachiocephalic artery (BCA); Ascending Aorta (AA); Distal Descending Aorta (DDA); Left Renal Artery (LRA); and Infrarenal Aorta (IA). For each of these flow measurements mean and pulse (Max – Min) values were extracted for each snapshot. For the remaining branches of the computational model without a flow measurement, mean flow target values were obtained using the following assumptions:

**Figure 8:**
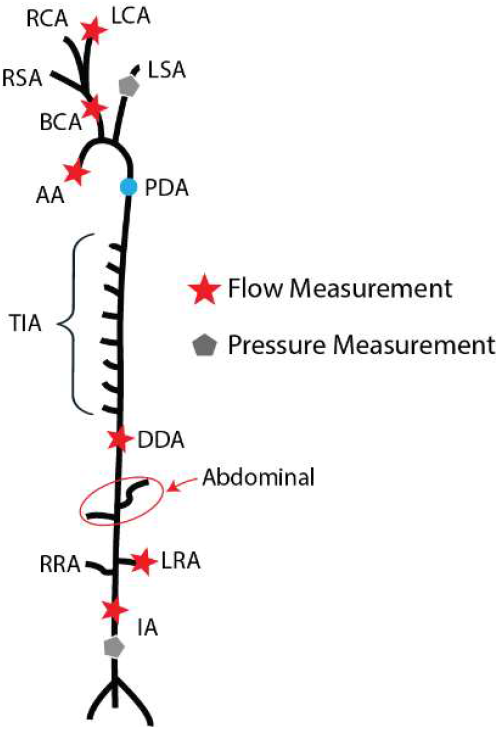
Simplified Flow and Pressure Probe locations.

**Figure 9:**
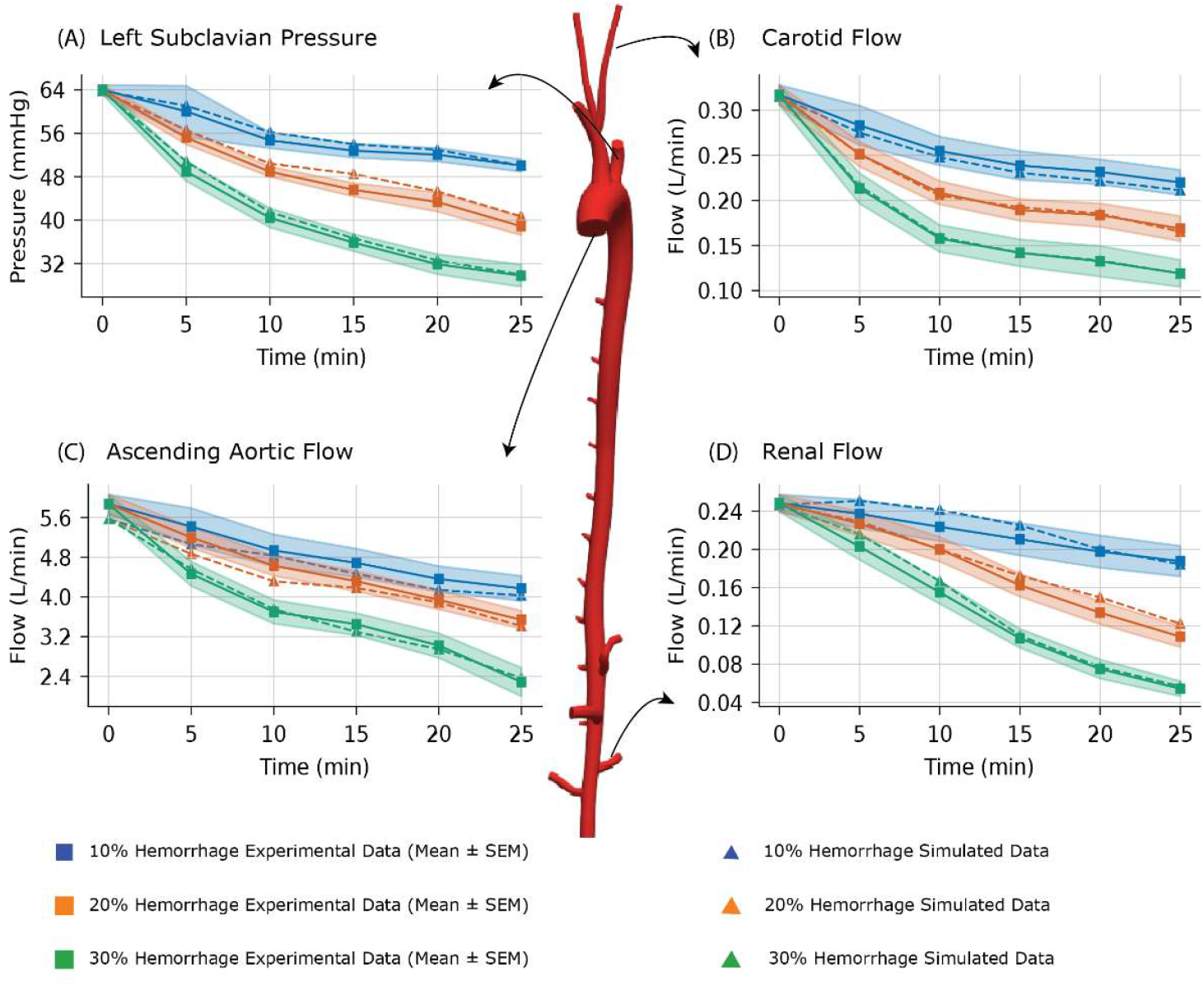
Representative group-level experimental (squares) and simulated (triangles) mean with shaded bands indicating ± SEM at each snapshot for (A) left subclavian pressure, (B) carotid flow, (C) ascending aortic flow, and (D) renal flow.

- Flow splits were assigned proportional to the cross-sectional area of each branch.
- Right carotid artery (RCA) mean flow was assumed to be the same as the measured LCA flow, scaled by the ratio of areas.
- Right renal artery (RRA) mean flow was assumed to be the same as the measured LRA flow, scaled by the ratio of areas.
- Measured BCA mean flow was split between the two carotid and right subclavian branches proportional to their areas.
- Mean left subclavian artery (LSA) flow was assumed to be the same as the estimated RSA artery flow, scaled by the ratio of areas.
- Mean proximal descending aorta (PDA) flow was computed by subtracting the measured BCA and estimated LSA flows from the AA flow.
- Total intercostal arteries (TIA) mean flow was estimated as the difference between the computed PDA mean flow and the measured DDA mean flow.
- Abdominal flow was defined as the combined flow to the celiac artery and superior mesenteric artery. Its mean value was estimated by subtracting LRA, RRA and IA mean flows from the measured DDA mean flow.

Arterial pressure was measured at the LSA and the IA (**Figure 8**). For each location, mean and pulse (Max – Min) pressure values were set as calibration targets for each snapshot.

## 3 Results

In the Results section, we first present a comparison between the experimental targets and the simulated results at a representative set of vascular locations to demonstrate the ability of the calibrated vascular model to reproduce the measured data. We then report the time history of a subset of vascular parameters, which provides branch-specific insight into how the inferred hemodynamic responses differ over the course of hemorrhage for the different groups. Lastly, we present a direct comparison between measured representative and simulated waveforms.

### 3.1 Experimental Group Level Means vs Simulation

In this section we compare mean hemodynamic values obtained with the calibrated vascular model with corresponding experimental mean targets for a subset of signals. Experimental variability is reported as the standard error of the mean (SEM), 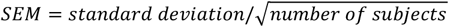 **Figure 6** shows comparisons for left subclavian pressure, ascending aortic flow, carotid flow, and renal flow.

Group-averaged experimental measurements showed a monotonic decline with both increasing hemorrhage severity and elapsed time (**Figure 9**). Mean left subclavian pressure dropped rapidly over the first 10 min (T0 - T10) in all groups, after which the 10% group partially stabilized, whereas the 20% and 30% groups continued to drift downward, with the 30% group exhibiting the largest overall reduction (**Figure 9A**). Ascending aortic flow showed a near-monotonic decline over time in all hemorrhage groups (**Figure 9C**). Regional flows displayed distinct dynamics: carotid flow fell sharply during the early phase (T0 - T10), particularly in the 20% and 30% groups, and then exhibited a slower decline thereafter (**Figure 9B**). In contrast, renal flow declined more gradually and continuously throughout the protocol without an obvious plateau (**Figure 9D**).

Across all cases, the calibrated vascular model reproduced the experimental group-mean trajectories and preserved the separation between hemorrhage groups, with simulated means generally remaining within the experimental uncertainty bands (SEM) at each snapshot.

### 3.2 Calibrated Parameter Trajectories Over Hemorrhage

The parameter trajectories shown in **Figure 10** are direct outputs of the vascular model calibration at each snapshot. The evolution of these parameters provides insights into distinct regional cardiovascular adaptations over hemorrhage (e.g., changes in regional resistance, effective arterial compliance, and venous volume recruitment) not directly measured in the experimental data.

**Figure 10:**
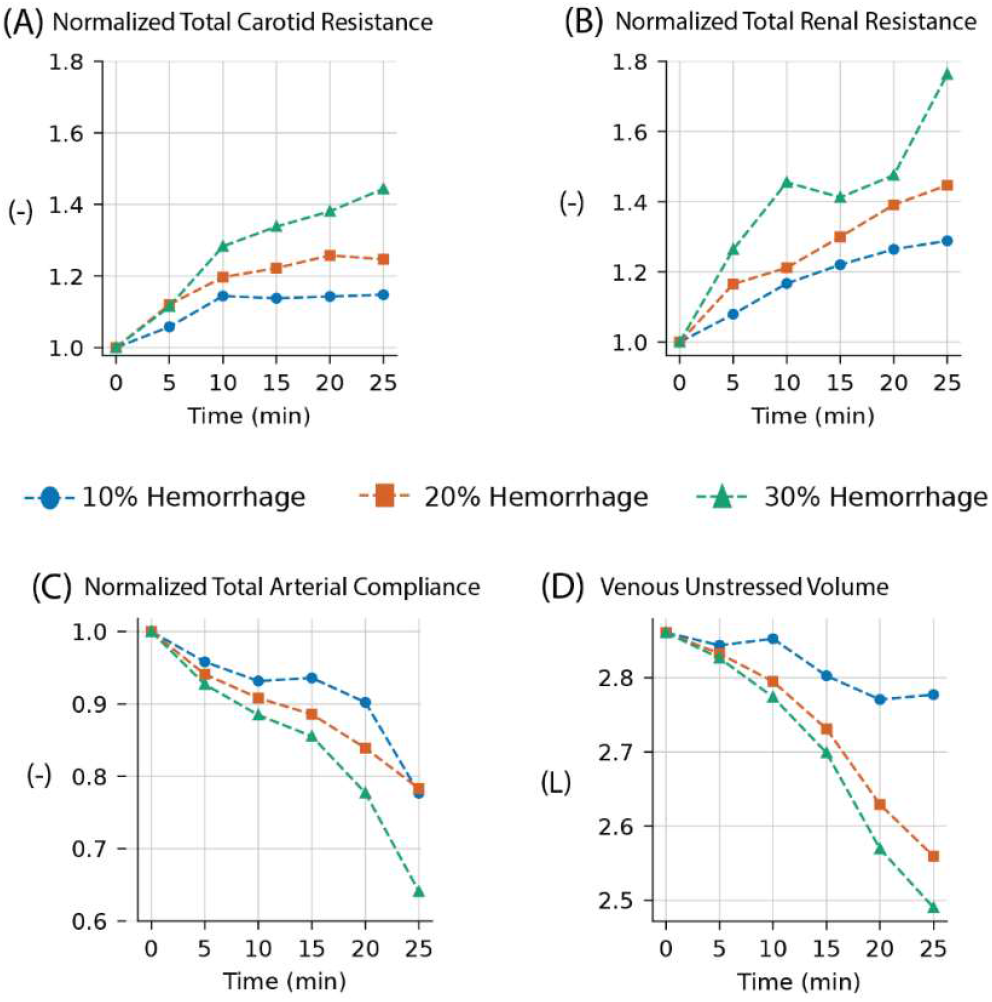
Evolution of calibrated vascular model parameters over hemorrhage: (A) normalized carotid total resistance, (B) normalized renal total resistance, (C) normalized total arterial compliance, and (D) venous unstressed blood volume *V*_*u*_.

Normalized total carotid resistance (defined as changes relative to calibrated values at baseline) increased over time in all hemorrhage groups, with clear separation by hemorrhage severity (**Figure 10A**). Divergence between groups became apparent after the T10-11 snapshot. The 10% group exhibited a plateauing, the 20% group showed modest increases in resistance, whereas the 30% group showed a continued rise through T25-26.

Normalized total renal resistance (**Figure 10B**) increased more rapidly than carotid resistance for all groups, with a pronounced early rise evident by T5-6 for the 30% hemorrhage group. Group separation appeared earlier in the renal bed compared to the carotid and persisted throughout hemorrhage. The 30% group exhibited a larger increase compared with the 10% and 20% groups.

Normalized total arterial compliance (**Figure 10C**) decreased with time in all hemorrhage groups. The 10% and 20% groups showed gradual declines, while the 30% group demonstrated an accelerated decrease after T10-11.

Venous unstressed volume *V*_*u*_ (**Figure 10D**) presented distinct patterns between the 10% and the more severe hemorrhage groups. *V*_*u*_ decreased modestly for the 10% group, showing a plateauing pattern after T15-16. In contrast, the 20% and 30% groups showed a progressive decreased in *V*_*u*_ over time, with the most severe drop (nearly 0.37 liters) in the 30% group. Table 2 compares changes in unstressed volume with total blood loss for each hemorrhage group.

**Table 2:** Changes in unstressed venous volume versus total blood loss for each hemorrhage group. An average initial blood volume of 3.8 liters was assumed for the average subject.

| Parameters | Hemorrhage Level |  |  |
| --- | --- | --- | --- |
|  | 10% | 20% | 30% |
| Total blood loss (L) | 0.38 | 0.76 | 1.15 |
| Total change in $V_u$ (L) | 0.1 | 0.3 | 0.37 |

### 3.3 Waveform Level Agreement at The Final Snapshot (T25-26)

A comparison between experimental and simulated waveforms at baseline (T0-1) and at the final snapshot (T25-26) for a subset of signals is presented in (**Figure 11**). This information provides insights into the agreement between data and simulation beyond simply the mean targets discussed earlier. Waveforms for LSA pressure, AA, LCA, and LRA, flows, as well as LV PV loops are presented.

**Figure 11:**
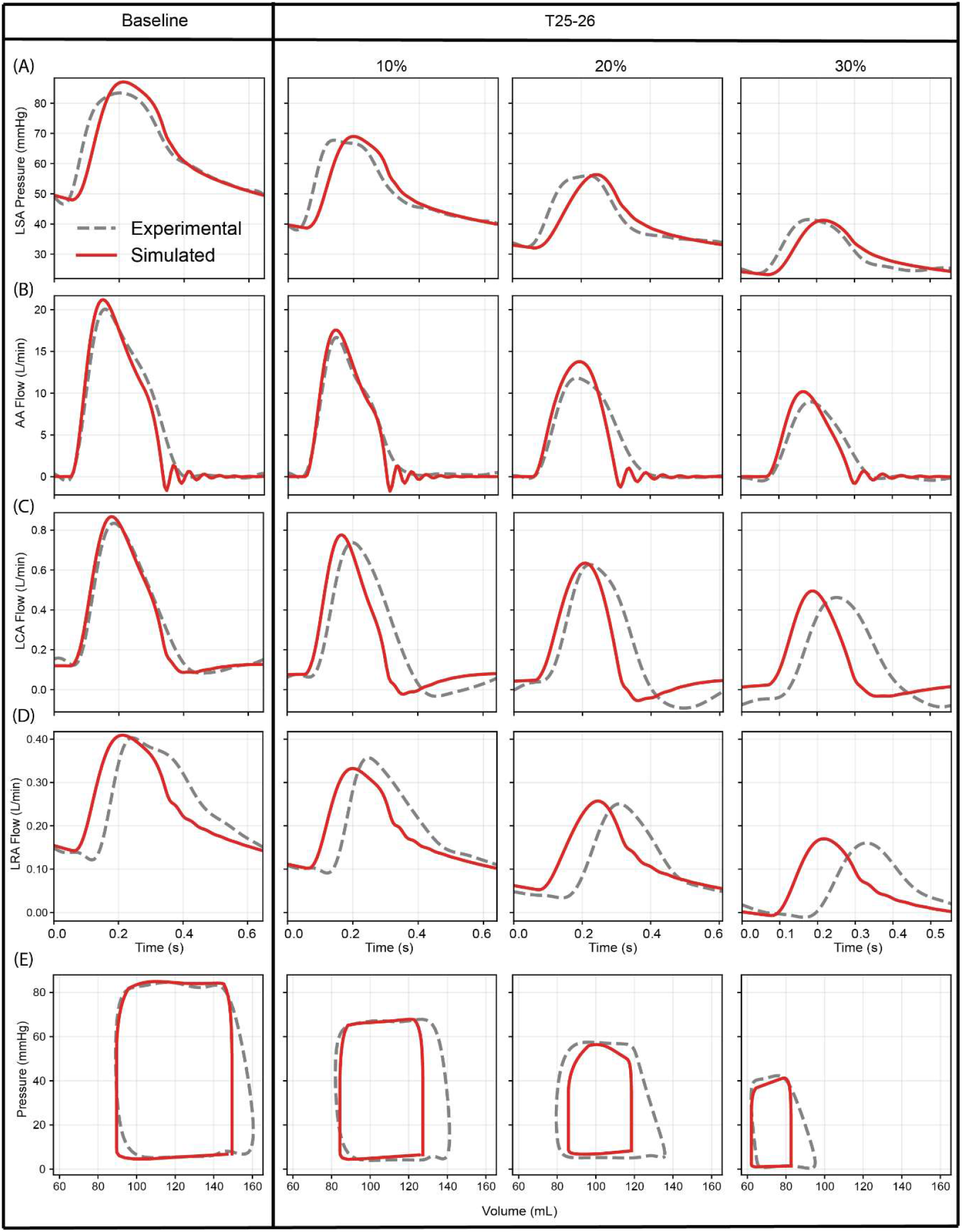
Representative waveform comparisons at baseline (T0-1) and final snapshot (T25–26) for 10%, 20%, and 30% hemorrhage. Experimental waveforms are shown as dashed gray traces and simulations as solid red traces for (**A**) left subclavian pressure, (**B**) ascending aortic flow, (**C**) carotid flow, (**D**) renal flow, and (**E**) left-ventricular PV loops.

Simulated waveforms reproduced hemorrhage severity-dependent waveforms in both shape and pulsatile magnitude across all signals. Relative to baseline, the final snapshot (T25-26) waveforms show a progressive loss of pulsatility with increasing blood loss. Despite the relatively modest changes in heart rate seen earlier (cf. Figure 4), the 30% group shows a shorter ejection period and cardiac cycle length compared to the other groups.

Simulated LSA pressures capture the systolic upstroke and diastolic decay and reproduces the progressive reduction in pulse pressure across hemorrhage groups (**Figure 11A**). Simulated AA flow similarly captures the reduced peak and shorter ejection period observed experimentally at higher blood loss (**Figure 11B**). Simulated LCA and LRA flows reproduce the decline in peak and pulsatility from baseline through 30% hemorrhage at the last snapshot (**Figure 11C-D**). And the simulated LV PV loops reflect the reduced filling and decreased stroke volume observed experimentally (**Figure 11E**). Lastly, across all signals, a phase lag becomes evident between simulated and experimental waveforms. This lag is more noticeable for higher blood loss, particularly for carotid and renal flows (**Figure 11C–D**).

## 4. Discussion

The primary aim of this study was to develop and calibrate a closed-loop zero-dimensional (0D) computational model of porcine aortic hemodynamics during progressive hemorrhage, to reproduce snapshot hemodynamics from controlled animal experiments. By integrating a physiologically interpretable model with detailed experimental data, the study provides mechanistic insight into the dynamic cardiovascular response to blood loss and lays the foundation for predictive modeling in trauma care settings.

### 4.1 Calibration Strategy and Model Identifiability

The calibration of the closed-loop lumped-parameter model (LPM) was carefully designed to balance model complexity with physiological interpretability. The modeling strategy restricts hemorrhage recalibration to a parsimonious set of parameters that are known for dominant short term actuator during hypovolemia. Our choice of calibrating just the peripheral Windkessel parameters (*R*_*p*_, *R*_*d*_, *C*) and venous unstressed volume *V*_*u*_ is consistent with other closed loop models in which acute arterial pressure control is achieved mainly through changes in vascular resistance and recruitment of venous unstressed volume. Ursino’s baroreflex model highlight an important role of venous unstressed volume in controlling the early response to acute blood loss [32]. Similarly, short term orthostatic stress models such as Heldt et al and Lau et al use a closed loop framework where rapid compensation is dominated by autonomically mediated changes in vascular tone and venous return, rather than slow structural remodeling [31], [36]. In practice, this restriction of parameters helped us obtain good agreement with the pressure–flow targets while keeping the parameter trajectories interpretable; however, it also means some physiological changes are absorbed into this reduced set. For instance, Dujardin and Stone [37], [38], showed in canine models that large-artery properties are altered by hemorrhage, which is not represented in our fixed-parameter approach we discuss the implications of this assumption below.

### 4.2 Mechanistic Interpretation of Calibrated Trajectories

Calibration produced trajectories for physiologically interpretable model parameters that provide a mechanistic explanation for the divergence between the hemorrhage groups. Agreement between simulated outputs and experimental targets is summarized in **Figure 9**, while **Figure 10** shows the evolution of calibrated total normalized carotid resistance, total normalized renal resistance, total normalized arterial compliance, and venous unstressed volume *V*_*u*_.

#### (1) Preload Limitation at Severe Hemorrhage

In the 30% hemorrhage group, an inflection is observed near T10–11, after which increasing peripheral resistance and declining venous unstressed volume *V*_*u*_ were no longer sufficient to preserve arterial pressure (**Figure 9-10**) [39]. This transition aligned with a breakdown in the ability to maintain cardiac output, consistent with established literature on hypovolemia, which describes a distinct progression from a compensated phase, maintained by vasoconstriction and venous unstressed volume *V*_*u*_ mobilization to a decompensatory phase marked by a drop in pressure once preload reserves are exhausted [40], [41].

#### (2) Regional Redistribution of Cardiac Output

Differential adaptation among vascular beds was evident. Renal resistance increased more steeply and earlier than carotid resistance, matching experimental observations in swine and underscoring a prioritized protection of cerebral circulation even as renal perfusion sharply declined (**Figure 9-10**). This pattern of differential regional vasoconstriction (renal > carotid) is physiologically consistent with experimental observations in swine showing pronounced reductions in renal perfusion during hemorrhagic hypotension, with relatively greater protection of perfusion to vital beds such as the cerebral circulation until compensation fails [42], [43], [44].

### 4.3 Model Efficacy Trade-Offs

A calibrated closed loop 0D framework is well suited for hemorrhage studies when the primary goals are to track systemic volume status, mean pressures, cardiac output, and regional redistribution over long periods of time, using parameters that remain physiologically interpretable. In a lumped-parameter (0D) formulation, pressures and flows are represented as functions of time without a spatial coordinate, which makes the model computationally efficient for real time clinical decision making [45]. However, without a spatial coordinate, a 0D model cannot reproduce finite pulse-wave transmission delays, reflection timing, or pulse-wave velocity effects. This limitation is apparent in our results as a progressive phase offset between simulated and experimental flow waveforms, most notably in distal beds such as the renal arteries. In the native circulation, pressure and flow waves travel at finite speed, and distal waveform timing depends on path length and arterial stiffness. The study acknowledges that future integration of 1D or spatially distributed models could address these limitations and improve waveform timing fidelity. Xiao and Alastruey have shown that one-dimensional (1D) networks introduce spatial propagation while being computationally efficient [46].

### 4.4 Limitations and Future Work

Several limitations define the current scope and motivate clear next steps.

#### Discrete snapshot calibration rather than a continuous control law

This study estimated parameters at discrete time points (“snapshots”) rather than embedding a continuously evolving autonomic control law. Snapshot calibration describes the system state at each one-minute interval, but it does not by itself model the between-snapshot trajectory without data. Closed-loop hemorrhage models that include explicit feedback mechanisms [15], [32] demonstrate how continuous control can be encoded within the model structure. Future work will therefore incorporate continuous reflex control including pressure regulation.

#### Anesthetized preparation and its effects

The experiments were performed in swine and under anesthesia, which can shift baseline hemodynamics and vascular tone. In particular, baseline MAP can be depressed under anesthesia, and this may place some vascular beds closer to their normal autoregulatory limit, complicating interpretation of hemodynamic transitions as purely hemorrhage-driven [47].

#### Fixed aortic properties and a path to address it

In this study, the proximal aortic RLC elements were fixed after the baseline calibration. This is a simplifying assumption, because experimental studies show that hemorrhage can increase proximal aortic characteristic impedance and that this change is attributed largely to altered aortic smooth muscle activity rather than to mean pressure alone. Dujardin and colleagues reported an increase in characteristic impedance after hemorrhage and concluded that changes in aortic smooth muscle tone were the primary driver. Stone and colleagues similarly demonstrated that volume changes modulate aortic characteristic impedance through smooth muscle activity [37], [38]. As a result, some hemorrhage-driven changes in pulsatile afterload may be represented in our model as effective changes in the peripheral Windkessel parameters rather than explicit proximal aortic adaptation. A practical extension is to incorporate a distributed arterial component (e.g., a 0D–1D hybrid), which would allow proximal aortic properties and wave transmission effects to be represented more directly while preserving the closed-loop calibration workflow.

#### Omissions of a coronary model

The present closed-loop circulation does not include an explicit coronary circulation. Coronary dynamics can be represented in lumped models, and dedicated coronary LPMs can represent the of coronary perfusion. Therefore, questions involving myocardial oxygen supply–demand matching or coronary perfusion would require extending the model.

#### Non-uniqueness of lumped-parameter calibration

Finally, inverse problems can be non-unique: different parameter combinations may reproduce similar outputs when the data are limited. This is a central point in inverse-problem physiology and motivates subset selection and identifiability-aware workflows. We reduced this risk by restricting parameters to physiologically plausible bounds and by calibrating a limited subset during hemorrhage, but non-uniqueness remains a structural limitation.

### 4.5 Conclusion

In conclusion, this study presents a data-driven, closed-loop zero-dimensional cardiovascular modeling framework calibrated directly to controlled hemorrhage experiments in swine, enabling reproduction of the time-evolving hemodynamic response across hemorrhage severities. The calibrated model captures the progression of pressure–volume loop contraction, mean arterial pressure, cardiac output, and regional flow redistribution. Beyond signal-level agreement, the inferred parameter trajectories provide mechanistic testable insight into adaptation during hemorrhage, including an increase in total peripheral resistance with regionally differentiated vasoconstriction and a substantial reduction in venous unstressed volume consistent with recruitment from the venous reservoir to support venous return under volume loss. Collectively, these results support the use of the proposed calibration framework as an interpretable in-silico framework for hypothesis testing during acute hemorrhage.

## 5. Acknowledgements

We’d like to acknowledge several students and staff who assisted with the animal experiments in this study at Wake Forest including Nate Bozeman, Jacob Dooley, Antonio Renaldo, and Hebah Soudan. Funding was provided by the National Institutes of Health Grant: R01-HL162633.

